# Predicting Attentional Focus: Heartbeat-Evoked Responses and Brain Dynamics During Interoceptive and Exteroceptive Information Processing

**DOI:** 10.1101/2023.11.03.565584

**Authors:** Emilia Fló, Laouen Belloli, Álvaro Cabana, Alessia Ruyan-Belabbas, Lise Jodaitis, Melanie Valente, Benjamin Rohaut, Lionel Naccache, Mario Rosanova, Angela Comanducci, Thomas Andrillon, Jacobo Sitt

## Abstract

Attention shapes our consciousness content and perception by increasing the probability of becoming aware and, or, better encode a selection of the incoming inner or outer sensory world. We designed a task to engage interoceptive and exteroceptive attention by orienting healthy participants to their heartbeats or auditory stimuli and investigated whether brain dynamics (Kolmogorov complexity – KC, permutation entropy – PE, weighted symbolic mutual information – wSMI, power spectrum density – PSD) and the heartbeat-evoked potential (HEP) distinguished interoceptive from exteroceptive covert attention. Exteroceptive attention yielded an overall flattening of the PSD, whereas during interoceptive attention there was a decrease in complexity, an increase in frontal connectivity and oscillations in the theta range, and a modulation of the HEP. Subject-level classifiers based on HEP features classified the attentional state of 17/20 participants. KC, PE, and wSMI showed comparable accuracy in classifying exteroceptive-interoceptive attention and exhibited a synergic behavior with the HEP features. PSD features demonstrated exceptional performance (20/20). Command-following was assessed in 5 brain-injured patients with a modified version of the task. An Unresponsive Wakefulness Syndrome/Vegetative State patient and a locked-in syndrome patient demonstrated a willful modulation of the HEP and the patient-level classifiers suggest that patients were complying with task instructions. Our findings show that directing attention to bodily rhythms or external stimuli elicits distinct neural responses that can be employed to track covert attention at the individual level. Importantly, the brain markers studied in this work provide multiple layers to explore information processing in disorders of conscious patients.

**Significance statement:** We show that directing attention to bodily rhythms or external stimuli induces specific neural responses, allowing for individual-level tracking of covert attention in healthy participants. The time-locked and dynamical brain markers identified as relevant to distinguish between interoceptive-exteroceptive attention were further assessed in a small cohort of brain-injured patients as an EEG proxy of command-following. We show for the first time the willful modulation of the HEP in an UWS/VS patient and a locked-in syndrome patient, along with significant classification accuracies when combining the explored features. Importantly, the brain markers studied in this work provide multiple layers to explore information processing in disorders of conscious patients, and expand the framework of heart-brain interactions employed for diagnostic purposes.

## Introduction

The brain continuously monitors the bodily and environmental signals, and the interplay between interoception and exteroception determines whether a change in state comes from within or from outside, triggering and shaping appropriate allostatic and behavioral responses (1–3). Although visceral signals are typically diffuse and not accessible to our conscious experience, interoception is considered to have a decisive role in perception, homeostatic responses, and motivational behaviors (4–7). Recent research has demonstrated that bodily rhythms contribute to general brain dynamics impacting the processing of external information (8–10). Notably, target detection of visual (11–13), somatosensory (14, 15), and auditory (16) stimuli is sensitive to the cardiac cycle phase. Conversely, external information can directly affect bodily signals as shown by modulations of the interbeat interval in anticipation of sensory stimuli (17) and after informative feedback error (18, 19), as well as following violations of statistical regularities (20). Hence, external and internal signal processing are intertwined and are sensitive to the state of the body and the state of the brain.

### Interoceptive attention

Attention has been described as a general mechanism that increases the detection of a desired signal while suppressing the response to irrelevant stimuli (21, 22). The modulatory effect of attention on sensory processing has been consistently shown for top-down attention on sound (23–25), touch (26), vision (27, 28), olfaction (29), taste(30). Crucially, visceral information can not only be passively filtered by the brain but attention can be directed toward bodily signals in a process known as interoceptive attention (5) impacting its cortical representation, which has been evidenced during interoceptive attention to respiratory cycle (31), and to the heartbeat (3).

### The heartbeat-evoked potential

The heart has been the preferred candidate to assess the effects of interoceptive attention at the individual level. The heartbeat is triggered by a dynamical pacemaker that is modulated by efferent brain pathways and inform the brain by ascendant pathways (9, 32), of which subjects are normally unaware. Brain response to the heartbeats can be measured by averaging time-locked EEG activity to the ECG waveform R peak (33). The resulting heartbeat-evoked potential (HEP) is considered to reflect the cortical processing of heart activity with and without awareness. The amplitude of the HEP is modulated by directing attention to the heart (33–37) (see (38) for a review), correlates with interoceptive awareness, measured as the accuracy in heartbeat detection (37, 39), and decreases with sleep depth (40).

### Brain dynamics during interoception and exteroception

Multiple processes related to top-down attention such as resource allocation, dynamical focus, inhibition, and selection have been associated with cortical oscillations and their entrainment (41–45). While brain dynamics of sensory processing have been extensively studied in the context of top-down attention for various exteroceptive modalities, investigations specifically comparing the electrophysiological response during interoceptive attention to the heart and exteroceptive attention, remain limited. Two studies based on visual and heartbeat detection tasks reported a trade-off between the HEP amplitude and visually evoked potentials during interoceptive attention, accompanied by an increase in parieto-occipital alpha power (36, 46). In addition, using intracranial recordings García-Cordero et al. (47) showed an increase in high-frequency oscillations (35-110 Hz) in interoception-related cortical regions during heartbeat tapping, and an increase in lower frequencies (1-35 Hz) when participants were tapping following an external rhythm. Together, these results suggest that time-locked and ongoing brain dynamics during interoceptive and exteroceptive attention can provide information on whether attention is oriented towards the internal or the external world.

### Perceptual learning and attention

Casting attention to a specific sensory channel may also enhance the representation of a stimulus that individuals are not actively scanning or may even be unaware of. An interesting case is posited by the perceptual learning of statistical regularities in noise. In the auditory modality, it has been shown that cross-trial repetitions of identical white noise fragments can result in persistent memory formations as indexed by participant’s detection accuracy (48, 49), memory evoked potentials (MEPs) (50), and intertrial phase coherence (ITPC) in the delta band (50, 51). Although the random white noise snippets used in the noise-memory paradigm have no semantic information or salient spectral features, their encoding is long-lasting (48), and brain signatures of a mnemonic response are elicited even when the repetitions are unbeknownst to the participants (50, 52, 53). Evidence on the effects of attention on implicit learning of these random acoustic patterns suggests that diverted attention hinders perceptual learning of the repetitions (52). Accordingly, orienting attention towards bodily rhythms should result in a worse encoding of these inconspicuous regularities, reflecting interoceptive or exteroceptive attention despite stimuli being hidden and orthogonal to task demands.

### Covert attention in unresponsive patients

As specified above, cortical and cardiac responses to external and internal stimuli are influenced by attention, and could therefore be used to detect covert attention. Developing measures of covert attention at the individual level can have major clinical implications as it could be applied to improve the detection of command-following responses (54–56) in non-communicative patients, such as patients who suffer from disorders of consciousness (DoC) (57). Assessing the level of conscious awareness in these patients poses a significant challenge as expertise is required to differentiate between reflexes and volitional behavior (58) and overt responses may be impaired (59). Indeed, the distinction of unresponsive wakeful syndrome, also referred to as vegetative state (UWS/VS), characterized by arousal without purposeful responses, from the minimally conscious state (MCS), where signs of intentional behavior are occasionally present, leads to significant misdiagnosis rates (60). Moreover, some patients show a dissociation between behavior and brain response, referred to as cognitive motor dissociation (CMD) (61), where residual consciousness can only be detected by functional neuroimaging methods. Finally, patients with Locked-In Syndrome (LIS), who are conscious but cannot show responses due to severe paralysis of almost all voluntary muscles except the eyes, may initially be misdiagnosed as UWS/VS (62). In this clinical scenario, active paradigms that measure command-following but that do not demand motor or verbal responses are especially appropriate, and a positive result provides significant information on the level of consciousness (63). Crucially, certain theories posit that the foundation of the sense of self hinges on defining the limits between oneself and the outside (4, 64–66). Therefore, probing the ability to recognize external and internal signals would give essential information on the level of self-awareness of these individuals.

### This study

We propose a novel task based on sustained selective attention to compare the effects of interoceptive and exteroceptive attention on the encoding of heartbeats, and salient auditory targets, as well as their effect on the perceptual learning of inconspicuous repetitions of white noise. We hypothesized that a heightened awareness of bodily signals should elicit an increased brain response to internal rhythms and a decreased response to external stimuli, with an opposite pattern during exteroceptive attention. Directing attention to external stimuli or internal rhythms should be characterized by specific brain dynamics, and should influence the ability to learn regularities passively. Importantly, we elaborated this study with the underlying motivation of applying this paradigm to obtain insights about the nature of attentive processes, and therefore our main goal was to investigate the suitability of different cortical and bodily measurements as markers to predict attentional focus at the individual level. Finally, we probed the clinical potential of our task on a small cohort of brain-injured patients.

## Materials and Methods

### Healthy participants

Twenty-two healthy volunteers participated in the task (13 females, average age, 30.63 ± 3.39). All participants reported having normal hearing and had not been exposed to the stimuli before the experiment. The study was approved by the ethics committee of the Facultad de Psicología, Universidad de la República (Uruguay). All participants gave informed consent and were not awarded any economic or academic retribution, according to the nationally established guidelines (Decree N379/008).

### Experimental design

The task consisted of 64 trials of 31 s of continuous white noise with bursts of amplitude-modulated noise (AmN) at random times. At the beginning of each trial, participants were instructed binaurally to close their eyes and focus their attention on the white noise (32 trials) or their heartbeats (32 trials), and to count the number of AmN or heartbeats, respectively. At the end of each trial, participants were audio-visually instructed to open their eyes and report the number using the keypad. Participants were not given any specific instruction on how to count their heartbeats but were exhorted to not measure their pulse. Half of the trials had embedded snippets of repeated white noise within and between trials (RepRN) that were not disclosed to participants (**Fig. 1A**). At the beginning of the experiment an 8-trial practice took place to ensure that participants understood the task. The practice stimuli were not repeated during the task and the noise repetitions were different from the ones used in the experiment. Stimuli presentation was coded in Psychopy 3.2.0 (67), and audio files were played using the sound library PTB in Windows 7 together with the Focusrite Scarlett 4i4 USB audio interface. Stimuli were presented binaurally through Etymotic ER3C tubal insert earphones and sound amplitude was adjusted for each participant during practice trials. The trials were presented in a randomized fashion and the inter-trial interval was randomly varied between 5 – 10 s.

**Fig. 1.**
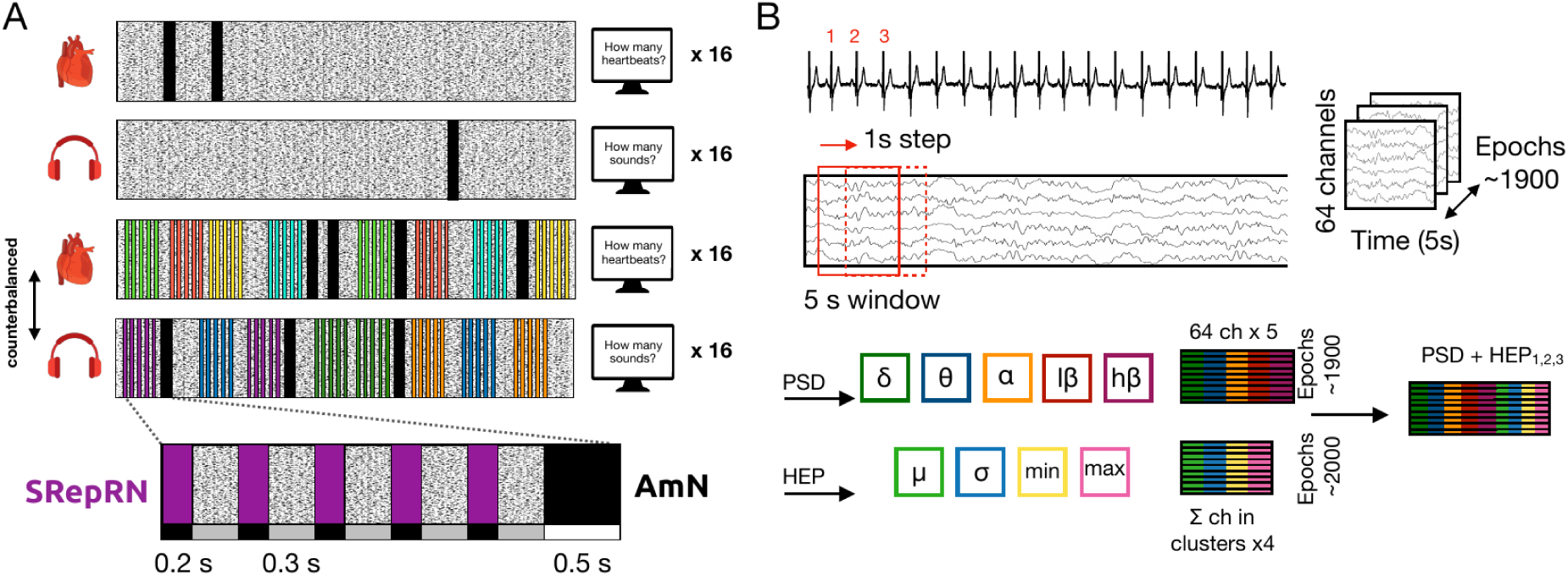
Experimental design and subject-level analysis. **A**. The task consisted of 64 trials of 31 s of continuous white noise with bursts of amplitude-modulated noise (AmN) at random times (represented with black lines over noise). On each trial, participants were instructed to focus their attention on the white noise (32 trials) or their heartbeats (32 trials) and were asked to report the counted number (AmN or heartbeats) at the end of each trial. Half of the trials had embedded snippets of repeated white noise within and between trials (SRepRN). Half of the participants were exposed to a set of four RepRN during the sound condition (represented with darker colored lines) and to another set of four RepRN during the heart attention condition (represented with lighter colored lines), this was counterbalanced across participants, such that each participant was exposed to a specific noise repetition in only one attentional condition. **B**. For the heartbeat-evoked potential (HEP) classifier, for each participant, a cluster permutation analysis was carried out to test for differences in HEP in the remaining participants. The channels and time points in the canonical clusters were used to extract the HEP features on the left-out subject data. For each electrode taking part in the clusters, the mean (μ), the standard deviation (std), the minimum (min), and the maximum (max) voltage in the time window spanning the cluster were extracted for each epoch of the subject withheld from the clustering analysis. For the PSD subject level classifier, each trial was segmented into 5 s sub-epochs with a sliding window of 1 s resulting in an overlap of 4 s between epochs. Power was obtained for each sub-epoch and averaged over the delta, theta, alpha, low-beta, and high-beta bands, resulting in five spectral features per channel. As an example of the combined classifiers, the PSD + HEP classifier combining both spectral and time-locked features is represented. The HEP features were derived from the average brain response to the heartbeats occurring within the 5-second sub-epoch, from which the spectral features were also extracted (i.e. heartbeats 1, 2, and 3 for the time window in red).

### Stimuli construction

AmN was constructed by multiplying 0.5 s of the running white noise background with a 40 Hz sinusoid at a modulation depth of 30%. For each trial, between 0 and 4 AmN were randomly included. Half of the trials (32 trials) included a concatenation of 5 copies of a structured noise repetition composed of a 0.2-s-long white noise snippet (RepRN) seamlessly concatenated to 0.3-s-long fresh noises. Eight different RepRN were created using different random seeds, such that four appeared during trials in which participants had to focus their attention on the sound (RepRN-Sound) and the other four RepRN were only included in trials of heart-directed attention (RepRN-Heart). The RepRNs assigned to each attentional condition were counterbalanced across participants. In each trial, the four RepRN concatenations were repeated twice, resulting in eight RepRN structured repetitions (SRepRN) within each trial (**Fig. 1A**).

### Physiological recordings and preprocessing

EEG and ECG signals were recorded using a Biosemi Active-Two system. Sixty-four Ag-AgCl scalp electrodes were placed on a head cap following the location and label of the 10-20 system, flat-type channels were placed on the left and right mastoid bones, and on the left and right collarbones to record cardiac activity. The signals were referenced online to the common mode sense (CMS, active electrode) and grounded to a passive electrode (Driven Right Leg, DRL). Data were digitized with a sample rate of 512 Hz with a fifth-order low-pass sinc filter with a –3 dB cutoff at 410Hz. As a backup, a second ECG recording was obtained following the same configuration but with a ground electrode positioned below the neck on the back of participants, and a respiration belt was used to record breathing. Both signals were recorded with a PowerLab 4/30 (ADInstruments) at a 400Hz sample rate. For two participants the backup ECG recording was used for the analyses. The analyses were conducted using MNE 1.0.3 (68).

### Interoceptive and exteroceptive accuracy

Interoceptive accuracy (IAcc) was measured for each trial according to the following equation. 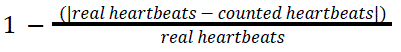. Values close to 1 indicate high similarity between reported heartbeats and actual heartbeats. Exteroceptive accuracy (EAcc) was defined as the percentage of AmN reported in relation to the total number of AmN presented across the entire experiment. The correlation between interoceptive and exteroceptive accuracy was evaluated using the Spearman correlation.

### EEG preprocessing

EEG data were processed according to the stimuli and the level of granularity of the analysis of interest, this resulted in 7 types of epochs: (1) For the heartbeat-evoked potential, data was filtered with a 30 Hz low-pass filter (one-pass zero-phase FIR filter with length 227 samples), epoched –0.5:0.8 s time-locked to the R peak of the ECG waveform, linearly detrended and referenced to the average of all channels. Epochs were not baseline corrected to avoid any contamination from the PQ component of the heartbeat. Heartbeats matching the moment of the AmN presentation were not included. (2) For the AmN, to remove alpha oscillations due to participants having their eyes closed, data was filtered with bandpass filter 0.2-7 Hz (one-pass zero-phase FIR filter with length 8449 samples), referenced to the average of all channels, epoched –0.1:0.85 s relative to sound onset, linearly detrended and baseline corrected 100 ms before sound onset. For the white noise repetitions, data was filtered following the preprocessing for the AmN but referenced to the average of the two mastoid electrodes. Subsequently, data underwent two types of epoching: (3) –0.4:3 s relative to sound onset thus comprising the 5 noise repetitions (SRepRN), and (4) from –0.05:0.5 s relative to each repeated noise (RepRN). For the ERP analysis of the SRepRN and for each RepRN, 0.2 s and 0.05 s to sound onset, were respectively used as a baseline. (5) In order to have a standard against which to compare the activity evoked by the noise repetitions, analog epochs to (3) and (4) were obtained from trials with plain white noise. This was carried out avoiding EEG data matching AmN presentation. (6) For the subject-level analyses, raw data was filtered with a bandpass filter 0.1 Hz – 30 Hz (one-pass zero-phase FIR filter with length 16897 samples), and referenced to the average of all channels. The interval between 1:30 s of each 31 s trial (to avoid onset/offset sound artifacts) was segmented into 5 s epochs with an overlap of 4 s. This procedure resulted in a comparable number of trials for the classifiers using ongoing brain activity as features, and HEP-based classifiers, as well as the combination of different families of features (see below). Finally, the 30 s epochs were segmented into (7) 5 s sub-epochs without overlap for group-level analyses of complexity, connectivity, and power. For all epoch types, autoreject 0.3.1 (69) was used to reject bad epochs and interpolate noisy channels. For all the analyses and participants, the number of observations was equalized between conditions taking into consideration temporal proximity and thus avoiding data imbalance. The ECG artifact (70) could not be completely removed using independent component analysis. Therefore we decided to keep the potential ECG contributions in the EEG but carry out analyses to check for potential differences in cardiac activity that could be driving our results. We tested for changes in heart-rate and heart-rate variability as well as differences in the ECG waveform across conditions.

### Power and intertrial coherence ERP

Time-frequency analyses were carried out on ERPs using the multitapers method to extract intertrial phase coherence (ITPC) and power in theta (1:4 Hz) and delta (4:8 Hz) for the heart time-locked epochs, and for 1:5 Hz for the SRepRN, with a 0.2 Hz resolution. The number of cycles used for each taper was set to two for the SRepRN analysis and to one for the HEP, with four as the time-half bandwidth of the multitapers. For the HEP power analysis decibel baseline conversion was carried on the subjects’ average power using the interval –300:-150 ms relative to the R peak, and for ITPC the mean phase value of that interval was used as a baseline (71).

### Group-level EEG analysis

Cluster-based permutation analyses were used to assess differences between attentional conditions for the AmN, the HEP, RepRN, and the SRepRN. For the noise repetitions, comparisons between the brain response to the noise repetitions and to plain white noise during the same attentional condition were also carried out. The time points tested for the HEP were –0.1:0.6 s, for the AmN the entire epoch was evaluated, for the RepRN the time interval 0:0.5 s was analyzed and for the SRepRN the time window evaluated was –0.3:3 s. Differences between conditions for all channels in the specified time windows were analyzed as follows: i) subject average for sound attention trials is subtracted from the subject average for heart attention trials (or the subject average response to RepRN during sound/heart attention trials is subtracted from the subject average to plain white noise during sound/heart attention condition), ii) a one-sample t-test is performed on every sample, iii) t values that exceed a dependent samples t-test threshold corresponding to an alpha level (p-value) of 0.025 (two-tailed, number of observations = 20) are clustered according to temporal and spatial proximity. The adjacency matrix for a Biosemi 64 channel layout as defined in Fieldtrip was used and 1 was the maximum distance between samples to be considered temporally connected. iv) t values for each data point within each cluster are summed to obtain a summed t statistic per cluster (t-sum), v) 2000 permutations of the data are computed, and for each permutation, the cluster with the biggest-summed t statistic is kept to obtain a null hypothesis distribution, vi) the proportion of clusters from the null hypothesis with more extreme values than the cluster obtained from the observed data yields the p-value for a given cluster. For the AmN the critical alpha level was set to 0.01 to avoid one extreme cluster. The same analyses were carried out for power and phase obtained from the HEP (the time window evaluated was –0.3:0.6 s) by averaging over the delta, theta, and alpha frequency bands. Likewise, for the SRepN we evaluated ITPC over the –0.3:3 s interval by averaging the ITPC values across 1:5 Hz. Cluster permutation analyses were conducted for subject-average periodic and aperiodic spectral components, complexity, and connectivity measures obtained from the non-overlapping 5 s epochs following the methods described below. The level of significance was established at ɑ = 0.05.

### Aperiodic and oscillatory dynamics

The FOOOF 1.0 library (72) was used to model the subject-average power spectrums for the frequency range 1:30 Hz to properly characterize the ongoing rhythmic and aperiodic activity during both attentional conditions. Model fitting was performed with the following parameters: aperiodic mode was set to ‘fixed’, peak width limits = [1,5], the maximum number of peaks = 4, peak threshold = 1.5, minimum peak height = 0.05. The overall average power spectrum showed peaks on the theta (4:8 Hz), alpha (8:14), and beta range (14:25 Hz). Therefore, for each model, we extracted the center frequency (CF), aperiodic-adjusted power (APW), and bandwidth (BW) for those oscillatory components. If for a given fit no oscillatory component was found, it was replaced with zero.

### Subject-level EEG analysis

In order to classify the attentional focus for each participant, evoked, connectivity, information theory, and spectral markers were extracted and fed to subject-level classifiers. The evoked features were computed from each brain response to the heart, and the rest of the features were extracted from the 5 s sub-epochs. Adaboosts classifiers with decision trees as base estimators (73) were implemented using Scikit-learn v1.0.2 (74). The number of decision trees was set to 1000 and the maximum depth of each decision tree was set to 1. The splitting criteria used for each decision tree was set to ‘gini’. To avoid data leakage, we followed a grouped stratified k-fold cross-validation procedure with 8 folds where on each fold all sub-epochs (or brain response to the heart) from the same trial were grouped either in the train or test data. For each classifier, the mean accuracy across folds, measured as the area under the receiver operating characteristic curve (AUC), was obtained and significance was evaluated using a non-parametric statistical approach (75). The labels for the observations were randomly permuted 500 times and for each permutation, the mean classifier accuracy was obtained. We compared the mean accuracy of our original data against the empirical null distribution of classification accuracies. The proportion of null classification accuracies greater than the AUC of the original data yielded our p-values. The level of significance was established at ɑ = 0.05. The resulting p-values were corrected using the false discovery rate method (76).

### HEP features

In order to obtain the HEP features a leave-one-out approach was implemented as follows. Cluster permutation analyses were carried out on all subjects except the subject for which the feature extraction and subsequent classification were going to be computed. For the canonical clusters obtained, clusters with p-values < 0.05 were selected. For each electrode taking part in the selected clusters, the mean, the standard deviation, the maximum, and the minimum voltage in the time window spanning the cluster were extracted for each epoch of the subject withheld from the clustering analysis. This resulted in 4 features per channel in the clusters. As a control, the same temporal windows were used to extract analog features from the ECG channel resulting in four features per cluster.

### Spectral, complexity, and connectivity features

For each subject, a Laplacian transformation (77) was applied on the channels of the 5 s sub-epochs to reduce the effect of volume conduction and obtain less correlated sensor signals. Power spectral density for frequencies between 1:30 Hz was obtained using multitapers as implemented in the psd_multitaper function in MNE. Power was averaged across delta (1:4 Hz), theta (4:8 Hz), alpha (8:14 Hz), low-beta (14:20 Hz), and high-beta (20:30 Hz) bands for each channel, resulting in 320 spectral features per sub-epoch.

To assess brain dynamics during interoceptive and exteroceptive attention, permutation entropy (PE), symbolic weighted mutual information (wSMI), and Kolmogorov complexity (KC) markers were computed. The selection of these complexity and connectivity markers was driven by previous work showing their efficacy in classifying conscious states in DoC patients (78, 79). Furthermore, these metrics capture non-linear dynamics and complement spectral and time-locked features. PE (80) quantifies the irregularity of the signal in each channel by examining the distribution of ordinal patterns within the data after transforming the EEG signal into symbolic representations. KC is another measure of complexity that assesses the predictability of the sensor signal by quantifying its compressibility. Finally, wSMI (81) estimates the connectivity between channel pairs by transforming the signal following the method for PE and computing the joint probability of the symbols in the signal. To avoid a disproportion between the number of observations and the number of features, for the wSMI classifier, we used only 32 channels in a 1020 configuration (496 features). These markers were obtained using the nice python library (78), and PE and wSMI were evaluated in the theta band which has been shown to provide a more accurate classification of patients and is the frequency band linked to the HEP generator (71).

### Time-locked and dynamical features

In order to evaluate a synergistic effect of the time-locked and dynamical features to classify attentional focus, combined classifiers were implemented joining the HEP features with each of the dynamical features (power, KC, PE, and wSMI). Brain responses to heartbeats arising during the 5 s sub-epochs were averaged, and from the evoked activity the HEP features (mean, std, min, max) were extracted as detailed above (**Fig 1. B**). If for a sub-epoch all concurrent heartbeats were discarded during epoch rejection, the median across all the observations was used to replace missing data.

### brain-injured patients

Seven brain-injured patients participated in the study, three patients were assessed at the IRCCS Santa Maria Nascente Fondazione Don Carlo Gnocchi ONLUS, Milan (Italy), and four patients at the Pitié-Salpêtrière Hospital, Paris (France) in the context of the EU-funded multicentric project Perbrain (82). Two Milan patients could not complete the task due to technical issues and were discarded from the analysis. The demographics and clinical information for the remaining five patients are listed in **Table S1**. All patients were in a subacute state (0.5:1.5 months since injury), except for M1 (∼8 months since injury), a chronic DoC patient. The assessment was performed following the ethical standards of the Helsinki Declaration (1964) and its later amendments and was approved by the local committees of each center (Comité de protection des personnes Ile de France I, #2013-A00106-39 and ethics committee section “IRCCS Fondazione Don Carlo Gnocchi” of ethics committee IRCCS Regione Lombardia, protocol number 32/2021/CE_FdG/FC/SA). Informed consent was obtained from the legal guardians of the patients before enrolling them in the study. The Coma Recovery Scale-Revised (CRS-R) (83) was performed by experienced neurologists on the same day as the task. All patients had behavioral responses to sound or cortical auditory responses assessed with the local-global paradigm (84).

## Results

We presented white noise with embedded salient auditory targets only, or with the same targets together with specific white noise repetitions to which participants were naive. Exteroceptive attention or interoceptive attention was elicited by asking participants at the beginning of each trial to report the number of targets or the number of heartbeats they felt. EEG, ECG, and respiration were recorded. Event-related potentials, oscillations, complexity, and connectivity were assessed at the group level, and subject-level classifiers were used together with time-locked and dynamical features to classify attentional focus.

### Task performance of healthy participants: heartbeats and AmN count

Healthy participants were able to focus their attention on the sound during sound attention trials, as shown by the proportion of AmN reported (**Fig. 2A**). The performance during interoceptive attention showed greater variability across participants but a correlation between interoceptive and exteroceptive accuracy was found (⍴(20) = 0.63, p = 0.002), suggesting an overall engagement in the task. In order to maximize brain responses to the noise repetitions, trials had a high density of RepRN, importantly, the RepRN did not interfere with participants’ ability to detect their own heartbeats, as interoceptive accuracy did not differ between trials with plain white noise and trials with embedded noise repetitions (ANOVA F(1,21) = 0.056, p-value = 0.82) (**SFig. 3**). The detection of AmN was effortless for participants, therefore the low performance of subjects 1 and 2 (<70%) was interpreted as a lack of engagement in the task, leading us to exclude them from further analyses.

**Fig. 2.**
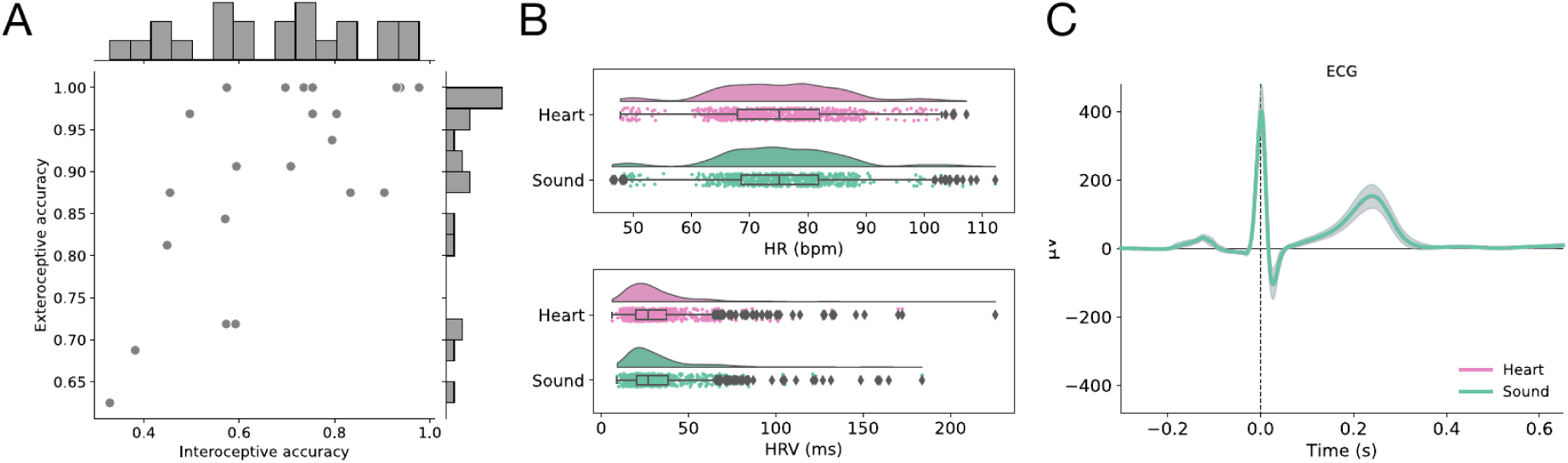
Task performance across subjects and heart activity. **A**. Correlation between exteroceptive accuracy as the percentage of AmN reported over the total number of AmN presented during trials of sound-directed attention) and the mean interoceptive accuracy across trials of heart-directed attention. Subjects 1 and 2 were discarded from further analysis due to low exteroceptive accuracy (<70%). **B**. Heart rate (top) and heart rate variability (down) for all trials during heart and sound-directed attention. **C**. Average ECG waveform for all subjects during sound and heart attention conditions with a 95% bootstrap confidence interval.

**Fig 3.**
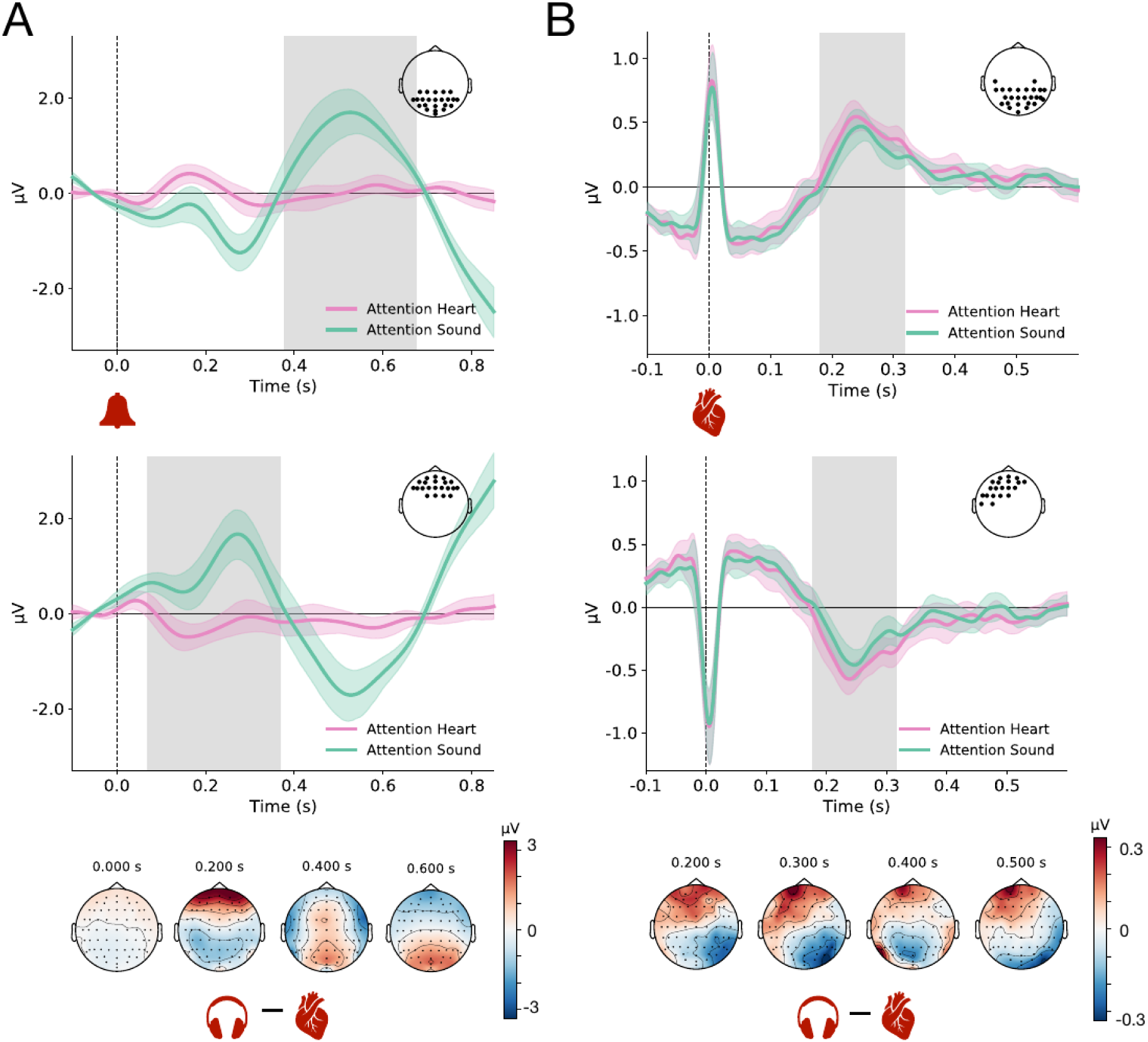
HEP and AmN evoked are oppositely modulated by interoceptive and exteroceptive attention. **A**. Brain response to amplitude-modulated noise. An early anterior (68 to 375 ms, t-sum = –9415, p-value < 0.001) and a later posterior (377 to 675 ms t-sum = –13983, p-value < 0.001) significant clusters were found. **B**. Heartbeat-evoked potential during heart and sound attention. Two significant clusters were obtained. A posterior cluster spanning from 179 to 318 ms (t-sum = 1382, p-value = 0.022) and an anterior cluster from 175 to 316 ms (t-sum = –1398, p-value = 0.022). The topography below each plot is the voltage difference between heart and sound conditions. Grey shading marks the temporal span of clusters and black dots indicate channels in clusters. The ERPs shown are the average voltage across channels for each cluster with a 95% bootstrap confidence interval.

### Heart activity and respiration are not modulated by exteroceptive-interoceptive attention

Mean Heart rate (HR) and mean heart rate variability (HRV) were measured for each 31 s trial and were contrasted during heart and sound attentional conditions (see supplementary materials). There were no differences in heart rate (HR heart = 75.13 BPM, HR sound = 75.44 BPM, t = 0.94, p = 0.36, β = 0.26, 95% CrI = [-0.43,0.96], BF = 0.14), nor in heart rate variability (HRV heart = 32.88 ms, HRV sound = 33.15 ms, t = 0.32, p = 0.75, β = 0.39, 95% CrI = [-1.38,2.17], BF = 0.12) between conditions (**Fig. 2B**). In order to assess whether the AmN prompted a cardiac deceleration, the interbeat interval for the first, second and third heartbeat following sound onset were compared to baseline (see supplementary materials). Independently of condition, there was no effect of AmN presentation on the interbeat intervals post target (ΔIBIB0 = 2.13, t = 0.85, p-value = 0.39, ΔIBIB1 = 0.67, t = 0.27, p-value = 0.79, ΔIBIB2 = –1.88, t = –0.75, p-value = 0.45) (**SFig. 1B**). Moreover, attention did not affect the ECG waveform as suggested by the negative results obtained for a point by point analysis (**SFig. 1A**) as well as a by a temporal cluster analysis (minimum cluster p-value = 0.46). Furthermore, we tested whether our task prompted changes in respiratory activity which could influence brain response to internal and external signals. Respiratory frequency did not differ across conditions (BR heart = 0.249 Hz, BR sound = 0.247 Hz, t = –0.61, p-value = 0.54), nor were there differences in the coefficient of variation of the breathing rate (CVBR heart = 0.174, CVBR sound = 0.166, t = –1.28, p-value = 0.20) (**SFig. 3**).

### HEP and AmN evoked responses are oppositely modulated by interoceptive and exteroceptive attention

The effects of attention on the cortical response to the heartbeats (HEP) was evaluated with a cluster permutation analysis. The analysis revealed two significant clusters. A posterior cluster comprised of 26 channels spanning from 179 to 318 ms (t-sum = 1382, p-value = 0.022) such that voltage was higher during heart-attention condition, and an anterior cluster comprised of 20 electrodes from 175:316 ms (t-sum = –1398, p-value = 0.022), for which voltage was more negative when attention was directed towards the heart (**Fig. 3B**). The difference in amplitude for the HEP did not correlate with subject’s interoceptive accuracy, and not all participants showed this modulation (**SFig. 4AB**). An increase in ITPC for the delta band was observed in posterior electrodes (t-sum = 7480, p-value < 0.001, time = –25:600 ms) (**SFig. 4C**), and for ITPC in the theta band (t-sum = –1381, p-value = 0.049, time = 344:600 ms). Moreover, no differences in power were obtained for the frequency bands tested, suggesting that the changes in the HEP are a result of phase modulations at low frequencies. Differences in brain response to amplitude-modulated noise targets (AmN) during the attentional conditions were also assessed using cluster permutation analysis. An increased response to AmN was observed when attention was directed to the sound (**Fig. 3A**) as indexed by five significant clusters spanning from 68 to 850 ms. We report here two of the clusters that summarize the topography of the effect. A later posterior cluster from 377 to 675 ms (t-sum = –13983, p-value < 0.001) and an early anterior cluster spanning the interval between 68 and 367 ms (t-sum = –8184, p-value < 0.002).

**Fig 4.**
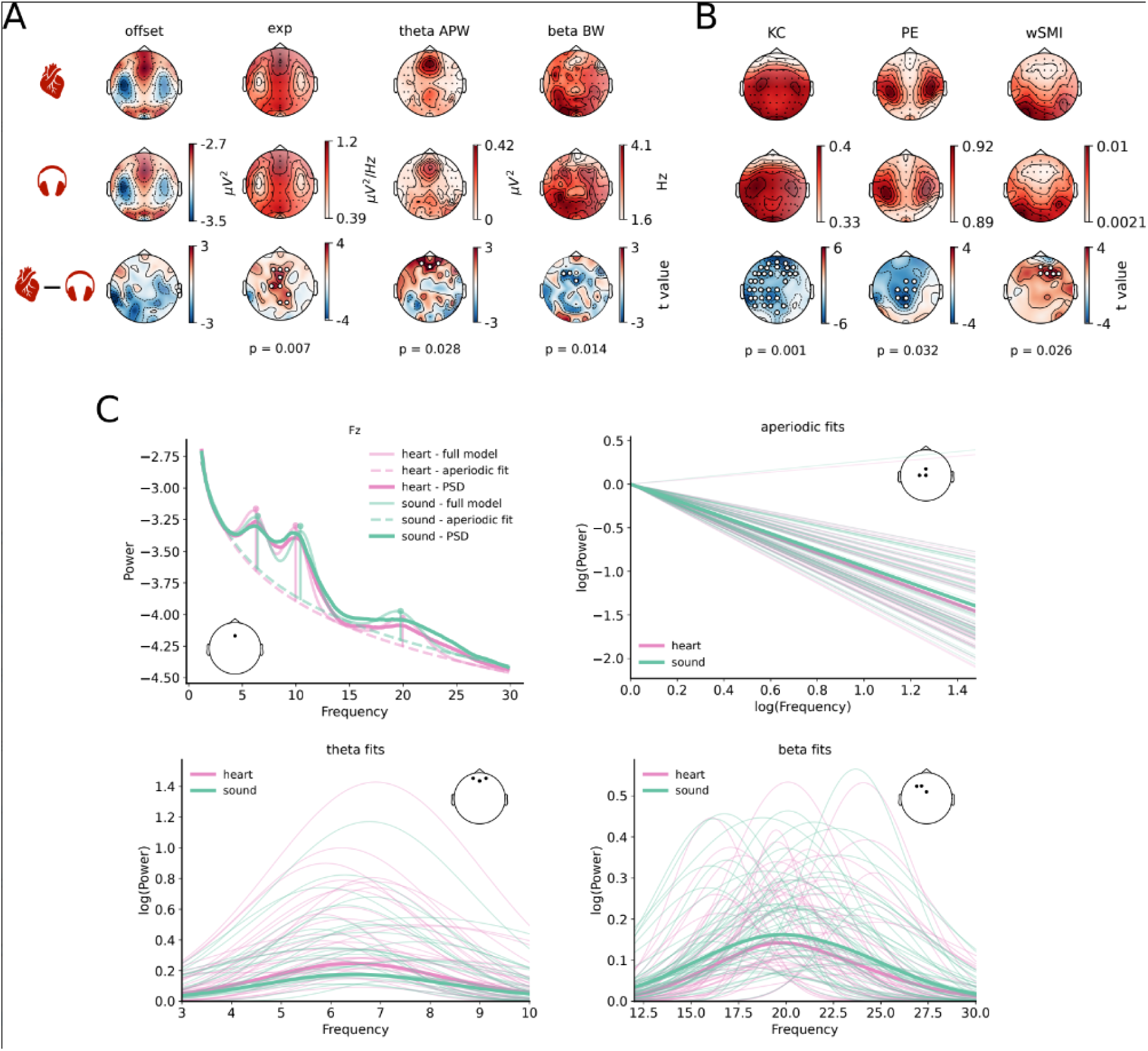
Interoceptive and exteroceptive attention show different brain dynamics. **A**. Periodic and aperiodic components of the power spectrum model fit during heart attention trials (top), sound attention trials (middle), and t-values for the difference between the components during both conditions (bottom). From left to right: offset value (off), aperiodic exponent (exp), and aperiodic-adjusted power (APW), and bandwidth (BW) for theta and beta bands. **B** Kolmogorov complexity (KC), permutation entropy (PE), weighted symbolic mutual information (wSMI). Top. Markers topography during attention to the heart. Middle. Markers topography during sound attention. Bottom. T-values for the difference between markers during heart attention and sound attention trials. White dots indicate sensors in significant clusters. **C**. Left top: average PSD for channel Fz with aperiodic and periodic model components for each condition. Right top: aperiodic fits for all subjects and conditions for Cz, C1, and FCz channels. Bottom left: periodic fits for the theta range (4:8 Hz) for all subjects and conditions for channels Fp1, Fp2, and AFz. Bottom right: periodic fits for the beta range (15:25 Hz) for all subjects and conditions for channels F3, FC1, and FCz.

### Interoceptive-exteroceptive attention and perceptual learning

In order to assess the effects of interoceptive and exteroceptive attention to inconspicuous noise repetitions, evoked responses to the repetitions (RepRN) and plain white noise (RN) were compared within each condition. Voltage differences were found between the RepRN and RN during sound attention as indexed by a widespread cluster (time = 0-209 ms. t-sum = 5586, p-value = 0.003). In addition, ITPC coherence in the delta band was higher during sound attention to the structured repetitions (SRepRN) compared to trials with plain white noise (time = 1.40-2.57 s, t-sum = 23072, p-value = 0.009) (**SFig. 5C**), and no difference in power was found (clusters p-values > 0.13). Conversely, no differences were obtained for the analog analyses between RepRN during trials of heart-directed attention and trials in which attention was directed to the heart but plain white noise was presented (**SFig. 5D**). In order to assess perceptual learning during heart and sound attention, evoked responses to the 1st to the 5th RepRN forming the SRepRN stimuli were separately averaged. Nevertheless, no positive results were obtained from the comparison of the 1st to 5th RepRN against plain white noise within each condition (**SFig. 5B**).

**Fig 5.**
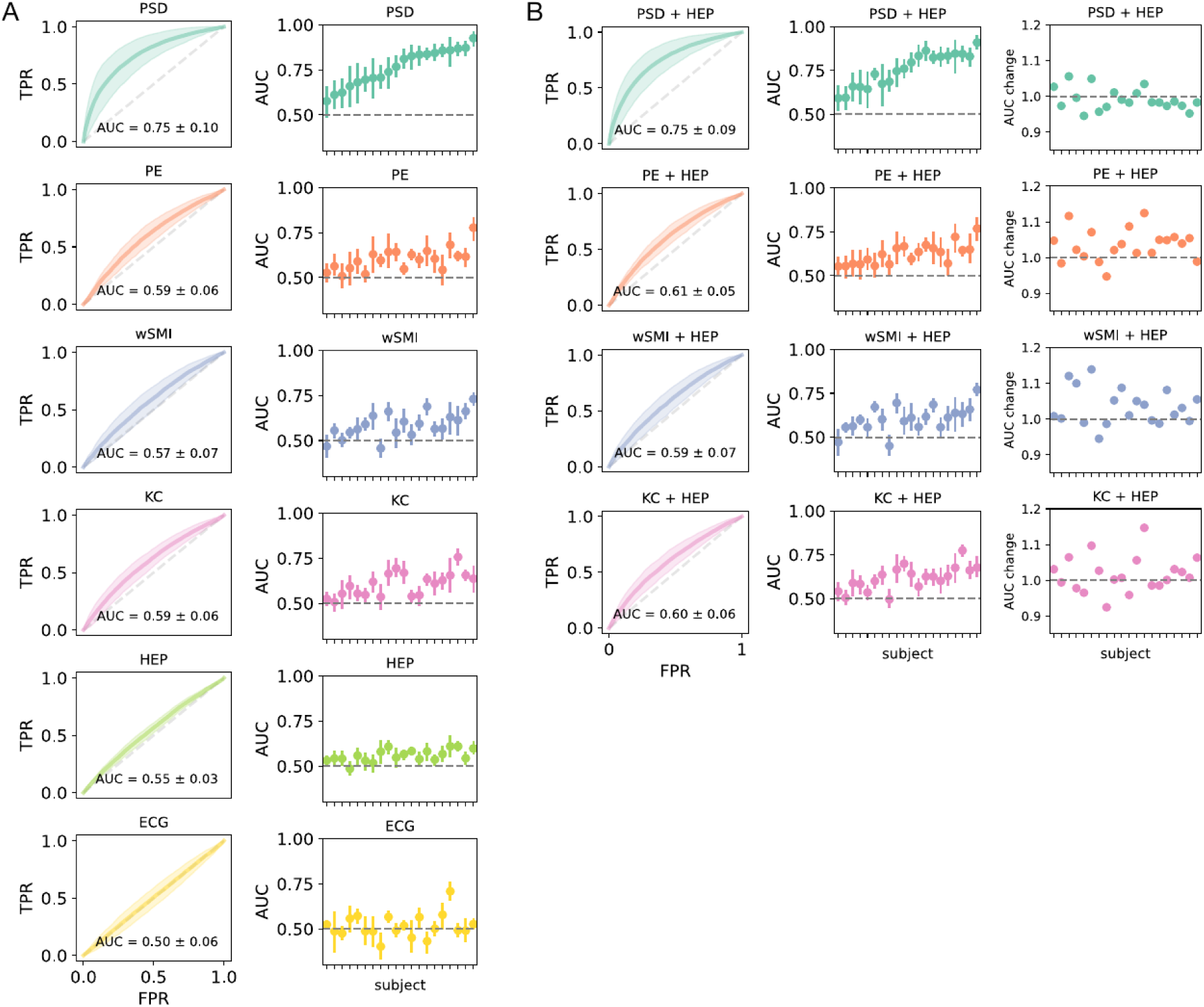
Brain dynamics and brain response to the heart are informative of attention orientation at the individual level. **A**. Subject-level classifiers for power spectral density (PSD), permutation entropy (PE), weighted symbolic mutual information (wSMI), Kolmogorov complexity (KC), heartbeat-evoked potential (HEP) and cardiac activity (ECG) features. Left. Average receiver operating characteristic (ROC) curves across cross-validation folds and subjects. Shading corresponds to the standard deviation. Right. Average AUC across folds with 95% bootstrap confidence interval for each subject and classifier. Subjects are sorted considering the AUC for the spectral density classifier. **B**. Combined classifier of dynamical features and HEP features. Left. Average receiver operating characteristic curves across cross-validation folds and subjects. Shading corresponds to the standard deviation. Middle. Average AUC across folds with 95% bootstrap confidence interval for each subject and classifier. Right. Ratio between the mean AUC for the individual classifier and the mean AUC for the combined classifier. Subjects are sorted considering the AUC for the spectral density classifier (not combined with HEP features)

### Rhythmic and aperiodic activity during exteroceptive-interoceptive attention

Brain oscillatory and aperiodic dynamics were contrasted between heart and sound-directed attention trials. A group effect of attentional condition was found for the aperiodic activity. Specifically, during interoceptive attention, the aperiodic exponent of the power spectrum was lower than during exteroceptive attention, and this effect was centrally localized in the scalp (tsum = 24.37, p-value = 0.007). In addition, differences were found in the oscillatory activity. The bandwidth of the beta band peak was smaller (tsum = –8.25, p-value = 0.014), and frontal theta power (tsum = 9.80, p-value = 0.028) was higher during heart-directed attention (**Fig. 4A, 4C**).

### Complexity and connectivity group analysis

Interoceptive and exteroceptive attention effects on brain dynamics were contrasted using Kolmogorov complexity, permutation entropy, and weighted symbolic mutual information. A significative difference was obtained for KC (t-sum = –112.77, p-value = 0.001) such that complexity was lower during interoceptive attention as shown by a widespread frontocentral parietal cluster. In addition, the analysis yielded differences in connectivity as indexed by a frontal cluster (t-sum = 17.87, p-value = 0.026) for which wSMI was higher during attention to the heart trials. Finally, permutation entropy was lower during interoceptive attention in centro-parietal electrodes (t-sum = –23.39, p-value = 0.032) (**Fig. 4B**).

### Time-locked activity, power, connectivity, and complexity classifiers

Adaboost subject-level classifiers were implemented using as features the time-locked and dynamical markers as well as integrated classifiers. Classifiers based on power across the delta, theta, alpha, low, and high-beta showed the best performance, accurately classifying all participants’ attentional condition with an overall AUC = 0.75 ± 0.10, with a performance superior to the one obtained for per-band classifiers (**SFig. 6**). The classifier based on the HEP features classified above chance 17 out of 20 participants with a mean AUC = 0.55 ± 0.03, and the AUC score was correlated to the number of epochs (⍴(18) = 0.46, p-value = 0.043) which was not the case for the PSD classifier (⍴(18) = 0.19, p-value = 0.41), suggesting that increasing the amount of data would yield more accurate results for HEP based classifiers. The ECG classifier only classified above chance 6 participants (AUC = 0.50 ± 0.06), which was unsurprising as no differences in cardiac activity were found in the group analyses. Complexity and connectivity-based classifiers showed comparable performance to the HEP-based classifier. The PE-based classifier was able to classify 17 participants (AUC = 0.59 ± 0.06), the wSMI-based classifier resulted in 16 participants being classified above chance (AUC = 0.57 ± 0.07) and the classifier based on Kolmogorov complexity accurately classified 18 participants (AUC = 0.59 ± 0.06) (**Fig. 5A**). No significant improvement in classification was obtained for the combination of spectral and HEP features (AUC = 0.75 ± 0.09, t = –1.48, df = 19, p-value = 0.92) nor when combined with KC features (AUC = 0.60 ± 0.06, t = 1.57, df = 19, p-value = 0.066). Nevertheless, an increase in overall classification was obtained when combining the time-locked features to PE (AUC = 0.61 ± 0.05, t = 3.63, df = 19, p-value < 0.001) and wSMI (AUC = 0.59 ± 0.07, t = 3.01, df = 19, p-value < 0.001) features, with three (total: 20 out of 20 participants) and two more participants (total: 18 out of 20 participants) being classified above chance, respectively (**Fig. 5B**).

**Fig 6.**
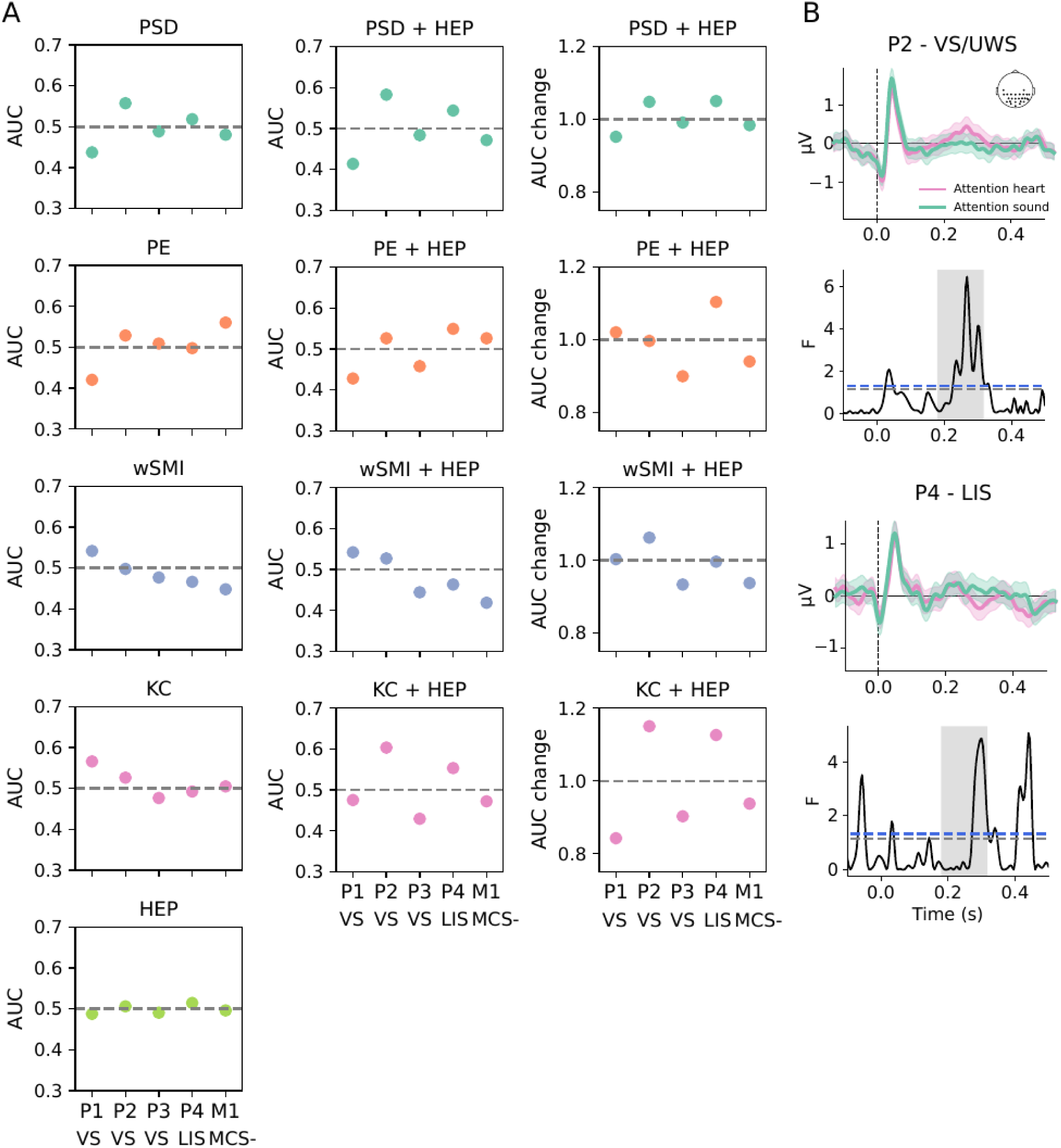
Brain dynamics and brain response to heartbeats to detect command following in non-communicative patients. **A**. Left: Subject-level classifiers for power spectral density (PSD), permutation entropy (PE), weighted symbolic mutual information (wSMI), Kolmogorov complexity (KC) and heartbeat-evoked potential (HEP) features. Average AUC across folds with 95% bootstrap confidence interval for each patient and classifier (VS: Unresponsive Wakefulness Syndrome/Vegetative State, LIS: Locked-in Syndrome, MCS-: Minimally conscious state minus. Middle: Combined classifier of dynamical features and HEP features. Average AUC across folds with 95% bootstrap confidence interval. Right: Ratio between the mean AUC for the individual classifier and the mean AUC for the combined classifier. **B**. HEP modulation by attention for patients P2 and P4. Top: average HEP response for electrodes for which an effect of attention is observed in healthy participants. Bottom. F values for point-by-point one-way ANOVA analysis between heartbeat-evoked responses during attention to heart and attention to sound. Black dotted lines represent p < 0.05. Blue dotted lines represent p < 0.05 after Bonferroni correction. Grey shading indicates the time window for the observed effect of attention on healthy participants.

### Brain-injured patients show a modulation of the HEP and ongoing activity consistent with command-following

Three UWS/VS, one MCS– and a LIS-diagnosed patients were presented with a modified version of the task (see supplementary materials for a description). Feature extraction was carried out and fed to classifiers to differentiate between trials of heart-directed or sound-directed attention as a proxy measure for command-following. For the LIS patient, we expected a brain response consistent with interoceptive and exteroceptive attention as these patients are conscious. Although, UWS/VS are patients who clinically do not show responses to the external world, and MCS-patients are individuals who show basic cortically mediated behaviors, such as visual fixation and pursuit and automatic responses (85, 86), in a misdiagnose scenario covert volitional responses could be present.

UWS/VS patient P1 was only classified by KC (AUC = 0.56 ± 0.03, p = 0.014, all rest p > 0.09). P2, also a UWS/VS patient, was classified above chance by the PSD features (AUC = 0.56 ± 0.03, p = 0.029), and although the HEP features were not sufficient for an above chance classification (AUC = 0.51 ± 0.01, p = 0.38), combining these time-locked features to the spectral and KC information improved the AUC scores (PSD: AUC = 0.58 ± 0.03, p = 0.002; KC: AUC = 0.60 ± 0.03, p = 0.002) yielding a successful classification. No EEG marker could differentiate between attentional conditions for UWS/VS patient P3. The attentional state of patient in LIS (P4) was not classified by the single marker classifiers, but combining the HEP features with the PSD, KC and PE resulted in an above chance classification (PSD+HEP: AUC = 0.54 ± 0.02, p = 0.042; KC+HEP: AUC = 0.55 ± 0.02, p = 0.013; PE+HEP: AUC = 0.55 ± 0.02, p = 0.026). Finally, MCS-patient tested at the Milan center (M1) was successfully classified only by the PE features (PE: AUC = 0.56 ± 0.02, p = 0.007) (**Fig. 6A**). Both patients who showed increased AUC scores when incorporating the HEP features (P2 and P4) show a modulation of this ERP on a time window and electrodes consistent with healthy participants response (**Fig. 6B**). A modulation of the HEP was not observed for the rest of the patients.

## Discussion

The aim of the present study was to develop and test an experimental paradigm that would allow to infer the attentional focus of an individual from brain and bodily responses to internal and external stimuli. The hypothesis underlying this work is that attention has an impact on our consciousness content and perception, such that it enhances the probability of becoming aware and, or, of better encoding a selection of the incoming inner or outer sensory world. For this purpose, we designed a task to engage interoceptive and exteroceptive attention by orienting participants to their heartbeats or to salient auditory stimuli, and measured their cortical, cardiac, and respiratory activity, while the effects of attention on passive encoding were probed using concealed noise repetitions.

### Interoceptive attention modulates cortical response to heartbeats

In agreement with previous findings (35, 47), our results show that directing attention to the heartbeat yields a modulation of the cortical response to the heartbeats. In addition to replicating the group effect on the HEP during interoceptive attention, we have shown the strong nature of this modulation, such that it is sufficient to accurately classify the attentional state at the single-subject level. Although HEP amplitude can be modulated by the respiratory cycle (87) exhalation and inhalation showed no differences between attentional conditions. Moreover, no modulation of the cardiac rhythm or the ECG waveform was found at the group level. In addition, cardiac activity was not effective in detecting covert attention. Together, our results suggest that the cortical differences were not driven by changes in the afferent signals but are a result of a top-down modulation on the cortical processing of the heartbeats consistent with predictive coding accounts (88–90). The HEP voltage modulation was accompanied by an increase in ITPC in the delta and theta band, with no differences in power, hinting at a phase-locking reset effect of attention on ongoing neural dynamics. It has been reported that increases in HEP amplitude are associated with increases in ITPC in these frequency bands (40, 71) and that during resting state the heartbeat induces a cortical synchronization in the theta band (91). In this line, we found an increase in connectivity in frontocentral electrodes during interoceptive attention as measured by wSMI for this low-frequency band.

The amplitude modulation of the HEP was not correlated to interoceptive accuracy scores. This is not surprising, as multiple criticisms for heartbeat counting as a measure of interoceptive awareness have been raised. Crucially, responses are influenced by the knowledge participants have of their own heart rates, and as these tasks are typically measured during rest, the heart rate variability is very small and therefore hard to perceive and report (92). Moreover, it has been reported that the belief about one’s own heart rate is a better predictor of the number counted than the actual quantity of heartbeats (93). Hence, the heightened encoding of this internal state could have occurred without participants being aware of their heartbeats. Furthermore, HEP modulation was variable across participants which may explain why some subjects were not successfully classified only with the HEP features. This intersubject variability is not surprising given the fact that individual differences in heartbeat awareness have been widely reported in the literature, and have been linked to body mass index (94), sex (95), age (96), emotion (97), and even sensory deprivation (98).

### Exteroceptive attention induces an overall gain in auditory processing

Brain response to AmN was substantially affected by attention. This was expected as participants had to detect an infrequent and salient sound, which typically elicits a P300 response (99). Although the effect of attention on perceptual learning of noise repetitions was not as prominent, the white noise repetitions were better encoded during sound attention trials, as shown by the ERP and ITPC results. No cortical response was found for the first repetition of the SRepRN, which would have indexed the long-term learning of the snippets and not a within SRepRN short-term learning effect. Our outcome could be the result of a lack of power, and increasing the number of repetition presentations in future work would clarify this. In fact, although an increased response is visually noticeable for the 3rd repetition during sound attention and the 4th repetition during heart attention in comparison to plain white noise (**SFig. 5B**), no statistical differences were obtained, supporting the hypothesis that we lacked power. From our results, we can only infer that during exteroceptive attention there was a discernable enhancement of auditory processing, leading to an overall increase in the brain response to the RepRN. Unlike previous research, our experimental design holds a distinctive advantage. Modulating attentional focus within the same task with all RepRN presented in both conditions across participants makes it less likely that results are driven by some seeds being more easily perceived than others and enables measuring concurrent learning effects.

### Brain dynamics during heartbeat and sound awareness

Multiple studies have focused on comparing the effects of internal attention (100) on the processing of external stimuli using paradigms based on mental operations such as mind wandering (101–105) or mental imagery (105), showing a decrease of sensory evoked potentials during attention to internal information, consistent with our findings. Concerning brain dynamics, these paradigms show increases in alpha power (106–108) associated with a top-down inhibition of cortical areas that would process distractor-relevant information (109), modulations of theta power reflecting working memory demands (110, 111) and increases in the theta-beta ratio during internal thought production and low-alertness (112, 113). However, scarce studies have directly compared brain dynamics during interoceptive and exteroceptive attention. In our study, and following previous research (47), power was lower during interoceptive attention compared to exteroceptive attention for frequencies between 1 and 30 Hz. Our methodological approach allowed us to attribute this difference to an overall change in aperiodic activity such that power at lower frequencies is increased in relation to power at higher frequencies during the interoceptive task. An overall steepening of PSD has been shown during anesthesia (114–116) as well as during sleep (116), and has mechanistically been attributed to changes in the excitation-inhibition ratio in the brain (115, 117). Functionally, increases in PSD slope have been observed during response inhibition (118, 119), and interpreted as a marker of top-down control required to sustain goal representations (120). During both interoceptive and exteroceptive attention participants were demanded to carry out a detection operation. Nevertheless, the heartbeats and the auditory stimuli we employed have intrinsically different properties and probably elicited different behaviors. Cardiac activity is a rhythmic stimulus, therefore counting heartbeats was a repetitive and sustained operation. On the contrary, AmNs occurred randomly and scarcely, and participants would incur in counting none or a few times on each trial. Finally, given the lack of salience of heartbeats, interoceptive-attention trials were probably more demanding for participants, which could explain the observed aperiodic activity differences. In addition, our results show that interoceptive attention was characterized by less complex or regular brain dynamics, with a topographical widespread pattern. Brain signal complexity is associated with the number of independent functional sources, such that the higher the complexity the less correlated the neural sources sustaining the overall activity (121). Lower complexity during interoceptive attention is consistent with a more overall stable brain configuration compared to the exteroceptive attention condition.

Together with changes in the background activity, the beta band peak was narrower, and theta power was higher during heart-attention trials. As mentioned, previous research has associated a higher theta-beta power ratio with episodes of mind wandering (122) which are typically elicited during sustained and repetitive tasks. Considering that the HEP was positively modulated by interoceptive attention, we argue that participants were actively engaged during the task making stimulus-independent thought unlikely, and the rhythmic changes observed would be a reflection of cognitive effort. In line with this, frontal theta has been associated with target detection (123), fatigue (124), and overall cognitive control (125).

PE and wSMI features combined with the HEP features enhanced the classification, signaling that they convey mutual information. This behavior was not observed in the case of spectral features or Kolmogorov complexity. For KC there was an improvement but it did not reach significance. Possibly the increase in information could not compensate for the detrimental effect of having a larger number of features in the resulting classifier. In the case of the spectral features, probably, the time-locked information is already embedded within the signal power decomposition, rendering both groups of features redundant and leading to no improvement. Including all spectral bands as features yielded a better classifier than using each power band by itself, which is consistent with an overall change in brain dynamics during our interoceptive-exteroceptive attention manipulation.

### HEP and dynamical features act synergically to classify the attentional state of two brain-injured patients

The potential of this tool to detect command-following was probed in a small group of brain-injured patients. Classification by the proposed features showed very different behaviors across patients which is expected in a small sample with such heterogenous etiologies. P1 showed inconsistent results across classifiers, with PE and PSD classifiers showing a below-chance performance, and KC an above-chance performance. This can occur when the models fit to noise in the training sets and is suggestive of no difference in neural activity during both attentional conditions. P3, behaviorally diagnosed as UWS/VS, did not show AUC values above chance for any of the markers, and M1, a MCS(-) patient, was only classified by the PE features, likely indexing no command-following behavior. Crucially, the brain responses of the LIS patient (P4) and a patient with a UWS/VS diagnosis (P2) were reliably classified by combining dynamical and HEP features. Although the classification accuracies were lower in these two patients, probably due to less sustained attention and a more noisy environment, the consistency across classifiers together with changes in the cortical responses to heartbeats topographically and temporally consistent with healthy participants’ responses, suggests that patients were complying with task instructions. The detection of command-following in the LIS represents a positive control of our task as consciousness is preserved in these patients and command-following is therefore expected. Importantly, our assessment suggests that P4 had higher levels of residual consciousness than conveyed behaviorally, and this would mean a mismatch between the patient’s conscious level and the clinical diagnosis. Our results expand the framework of heart-brain interactions employed for DoC diagnostic purposes (20, 126, 127), by showing for the first time the willful modulation of the HEP in two brain-injured patients with severe impairments of sensory, motor, and executive functions. Compared to other command-following tasks (54, 56), our paradigm possesses the advantage of contrasting brain responses to two active instructions (instead of active instructions versus resting state), which in addition are less demanding compared to executing complex imaginary behaviors or actual movements. Beyond command-following, we argue that the ability to distinguish between internal and external signals in DoC patients could be interpreted as a signature of preserved self-awareness. Moreover, the proposed task can provide information on different levels of information processing independently of the patient following the instructions, as passive cortical responses to sounds, and heartbeats can be measured, as well as assessing EEG markers that have already proven robust in indexing the state of consciousness in DoC patients (78, 79). Our results are a proof of concept of the potential of this novel tool to detect command-following among patients who are unable to convey explicit behavioral responses and the feasibility of its application in clinical settings. Future work assessing a bigger cohort of patients should be carried out to comprehensively evaluate its diagnostic as well as prognostic capabilities.

## Limitations

Some aspects of our study should be considered to improve future work. Introducing a resting state condition against which to compare the brain markers obtained during both types of attention would be a good strategy to allow a more mechanistic interpretation of the results. Furthermore, the implementation of trials with varying lengths would prevent participants from offering similar responses on a trial-by-trial basis on the number of heartbeats simply due to their expectation of a consistent heart rate. Another limitation is that an exhaustive analysis of the effects of interoceptive-exteroceptive attention on heart activity comprising frequency domain and non-linear measures could not be conducted as the length of trials was too short to provide reliable results (128, 129).

## Conclusions and future directions

In this work, we explored the modulatory effects of interoceptive-exteroceptive attention on the cortical processing of bodily and external signals. We report an overall gain in auditory processing during exteroceptive attention, as indexed by an increased cortical response to target sounds as well as a better encoding of noise repetitions, and a heightened cortical response for heartbeats during interoceptive attention. Exteroceptive attention was characterized by an overall power increase across the frequency range of 1-30 Hz, whereas during interoceptive attention there was a decrease in complexity, together with an increase in theta, and a decrease in beta oscillations. Our findings demonstrate that directing attention to bodily rhythms and the external world elicits distinct neural responses that can be employed to track covert attention at the individual level. Importantly, we show that the brain markers studied in this work can be useful in detecting EEG proxy of command-following in unresponsive patients. Crucially, the proposed paradigm provides multiple layers to explore information processing and awareness in these patients and requires equipment commonly available in clinical environments, rendering its application across centers straightforward.

## Declarations

### Ethics approval and consent to participate

The study was approved by the Comité de ética de la Facultad de Psicología, UdelaR, the comité de protection des personnes Ile de France I, #2013-A00106-39 and ethics committee section “IRCCS Fondazione Don Carlo Gnocchi” of ethics committee IRCCS Regione Lombardia, protocol number 32/2021/CE_FdG/FC/SA).

## Consent for publication

Not applicable

## Availability of data and materials

The authors declare that the data and code supporting the findings of this study are available upon reasonable request

## Authors’ contributions

EF analyzed data. EF collected the data. EF, LB, AC analyses conceptualization. EF, JS, TA study conceptualization and design. ARB, EF, LJ, MV, AC collected patients data. AC, MR, BR, LN clinical work, JS provided funding. EF wrote the manuscript. All authors have read and approved the manuscript

## Acknowledgments

The authors would like to thank the patients and close relatives as well as Micah G. Allen for his useful comments.

