## Supplementary material for "Predicting Attentional Focus: Heartbeat-Evoked Responses and Brain Dynamics During Interoceptive and Exteroceptive Information Processing"

*ECG analysis*

Raw data for the difference between the channel placed on the left and right collarbone was processed with Neurokit2 0.2.0 toolbox (Makowski et al., 2021). Data was filtered with a 0.5 Hz high-pass Butterworth filter (order = 5) and a 50 Hz Butterworth notch filter (order = 2). R peaks in each 31 s epoch were detected using the method 'neurokit'. Ectopic heartbeats were automatically identified and discarded and wrongly detected peaks were corrected by setting a specific minimum height for the R peak individually for each subject. Mean heart rate (HR) in beats per minute and heart rate variability as the root mean square of successive differences between normal heartbeats (RMSSD) were obtained for each epoch (Baek et al., 2015; Munoz et al., 2015; Salahuddin et al., 2007). For each participant, a value was considered an outlier and discarded if it was below or above 3 standard deviations. To test for differences between conditions, linear mixed-effects models were fitted to heart rate and heart rate variability with condition (heart - sound) as a fixed factor and subject as a random effect. Bayesian hierarchical models were implemented to quantify the probability that our data supported a difference in heart rate activity over a null model. Two analyses were conducted to test for differences in the ECG waveform between conditions. First, point-by-point two-tailed one-sample t-tests on the subjects' difference between the mean ECG waveform during attention to the sound and attention to the heart were conducted, and Bonferroni corrected for multiple comparisons. Secondly, a cluster permutation analysis was carried out by comparing the temporal cluster obtained for the observed t values against an empirical null distribution of clusters obtained for 2000 instances after randomly flipping the signs of the differences between conditions. The cluster-forming threshold value for a critical alpha of 0.025 was obtained for a t distribution with 20 observations (two subjects were discarded due to low performance, see below). Linear models were implemented in R Studio (77) with R version 3.6.3 (Team, 2017) using lme4 (Bates et al., 2015), rstan (Stan Development Team, 2023), and brms (Bürkner, 2017) libraries.

To test whether the sound presentation elicited a heart rate deceleration, heart rate variability was assessed by comparing the interbeat interval (IBI) for the first, second, and third heartbeat after AmN onset to the IBI for the beat immediately before the target presentation. The IBIs were referenced to the average IBI before the target onset (Skora et al., 2022). The resulting differences were z-scored and values were rejected if below or above 3 standard deviations. The effect of target presentation on heart rate variability was evaluated with a linear mixed-effect model contrasting the IBI for the different heartbeat positions with subject as a random factor.

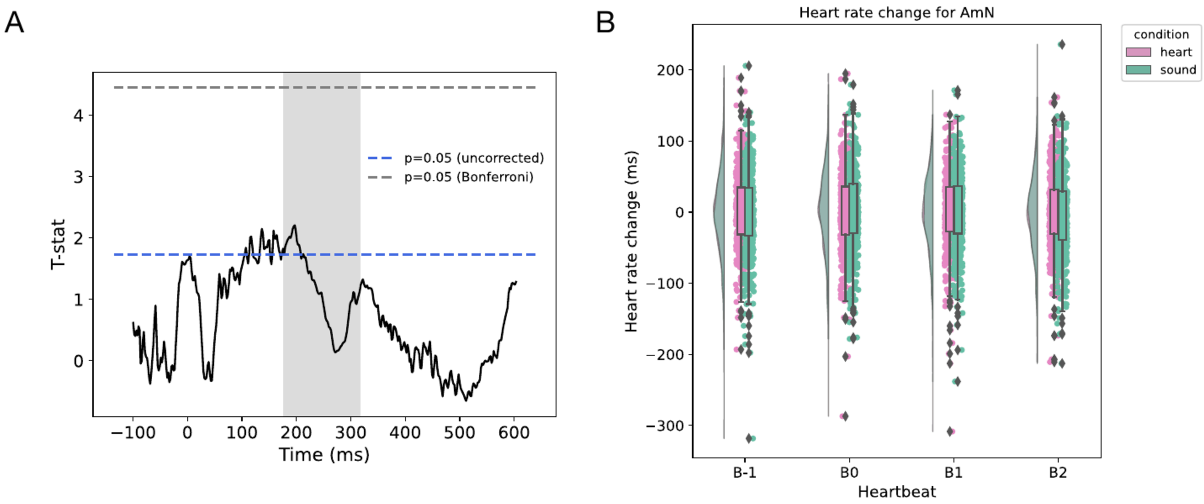


**SFig 1. A**. ECG waveform analysis. T-values were obtained from a two-tailed one-sample t-test on the subjects' difference between the mean ECG waveform during attention to the sound and during attention to the heart. The grey dotted line marks the threshold of significance after Bonferroni correction. Grey shading marks the HEP significant time points obtained from the cluster permutation analyses on the ERP. **B**. Target effect on cardiac activity. Heart rate change in relation to the average interbeat interval for heartbeat previous (B-1) to noise modulated in amplitude (AmN) onset. B0, B1, and B2 are the first, second, and third heartbeats after stimulus onset. No differences in ΔIBI were found between B0, B1, B2, and B-1.

*Respiration analysis*

Using breathmetrics toolbox (Noto et al., 2018) implemented in MATLAB 9.7 (2019b) the interbreath interval (IBrI), breathing rate (BR), and the coefficient of variation of the breathing rate (CVBR), measured as the standard deviation of the difference between inhale onsets divided by the average difference between inhale onset, were obtained. To test for differences between each of these independent variables and attentional condition, linear-mixed models were carried with condition (heart - sound) as a fixed factor and subject as a random effect.

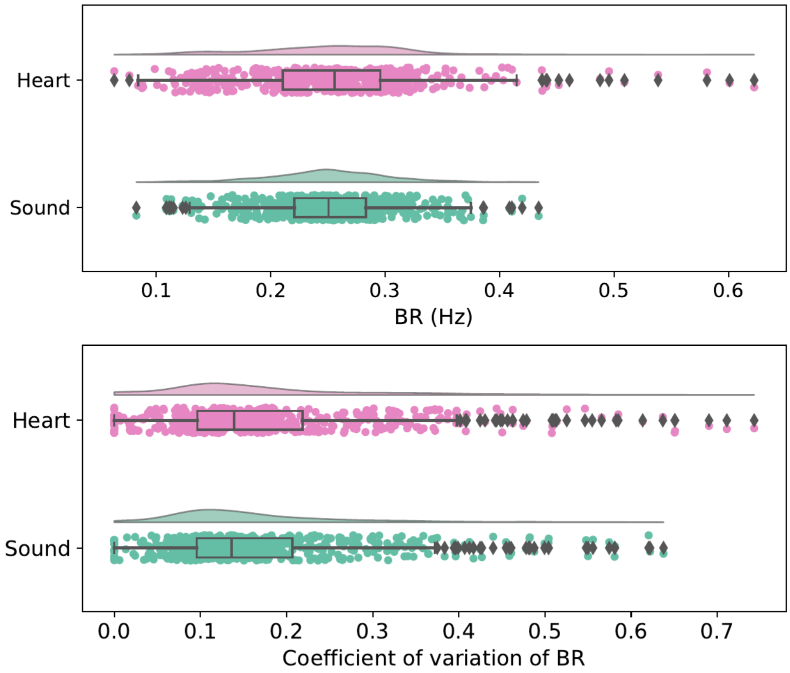


**SFig 2**. Attention effect on respiratory activity. Top: respiration frequency during heart-attention trials did not differ from sound-attention trials. Bottom: the coefficient of variation (standard deviation of the difference between inhale onsets over the average difference between inhale onsets) did not differ between attentional conditions.

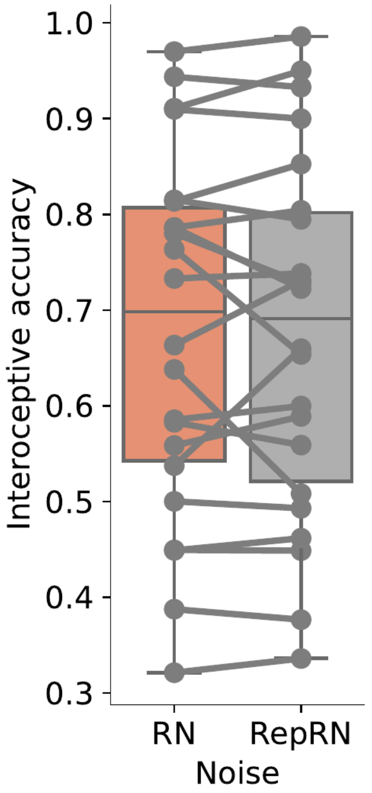


**SFig 3**. Noise repetitions and interoceptive accuracy. Mean interoceptive accuracy for each participant during rials with plain white noise (RN) and trials with embedded noise repetitions (RepRN).

*Correlation between HEP amplitude and interoceptive accuracy*

For each subject, the average voltage across channels and time points for each cluster resulting from the group analysis was correlated with their interoceptive accuracy using Spearman correlation. No correlation was found for the posterior cluster ($\rho(18) = 0.13$, p-value = 0.58) (Fig. S4A) nor for the anterior cluster ($\rho(18) = -0.03$, p-value = 0.89) (Fig. S4B).

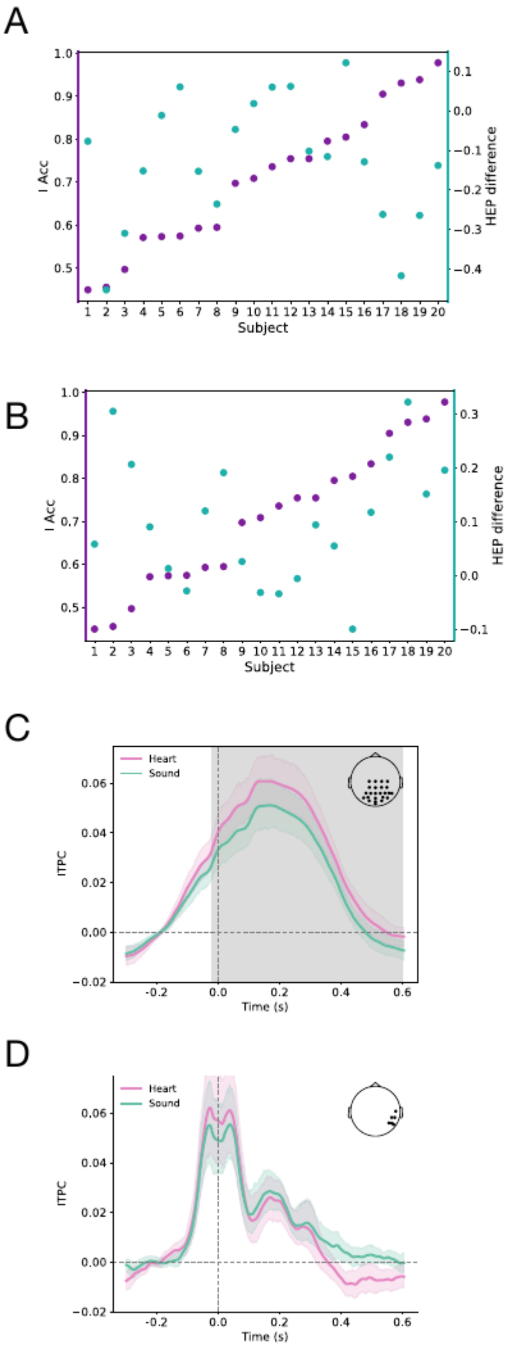


**SFig 4.** Correlation between mean amplitude for the time points and electrodes in the group level significant clusters for each subject and the mean interoceptive accuracy across the task. **A**. Mean amplitude for the time points and electrodes in the group level significant posterior cluster. **B**. Mean amplitude for the time points and electrodes in the group-level significant anterior cluster. **C**. Inter-trial phase coherence in the delta band for the heartbeat-evoked potential during attention to the heart (pink) and attention to the sound (green). Grey shading marks the temporal span of the significant cluster (t-sum = 7480, p-value < 0.001, time = -25:600 ms) and black dots indicate channels in clusters. **D**. Inter-trial phase coherence in the theta band for the heartbeat-evoked potential during attention to the heart (pink) and attention to the sound (green). Grey shading marks the temporal span of the significant cluster (t-sum = -1381, p-value = 0.049, time = 344:600 ms)) and black dots indicate channels in clusters.

The ERPs shown are the average voltage across channels for each cluster with a 95% bootstrap confidence interval.

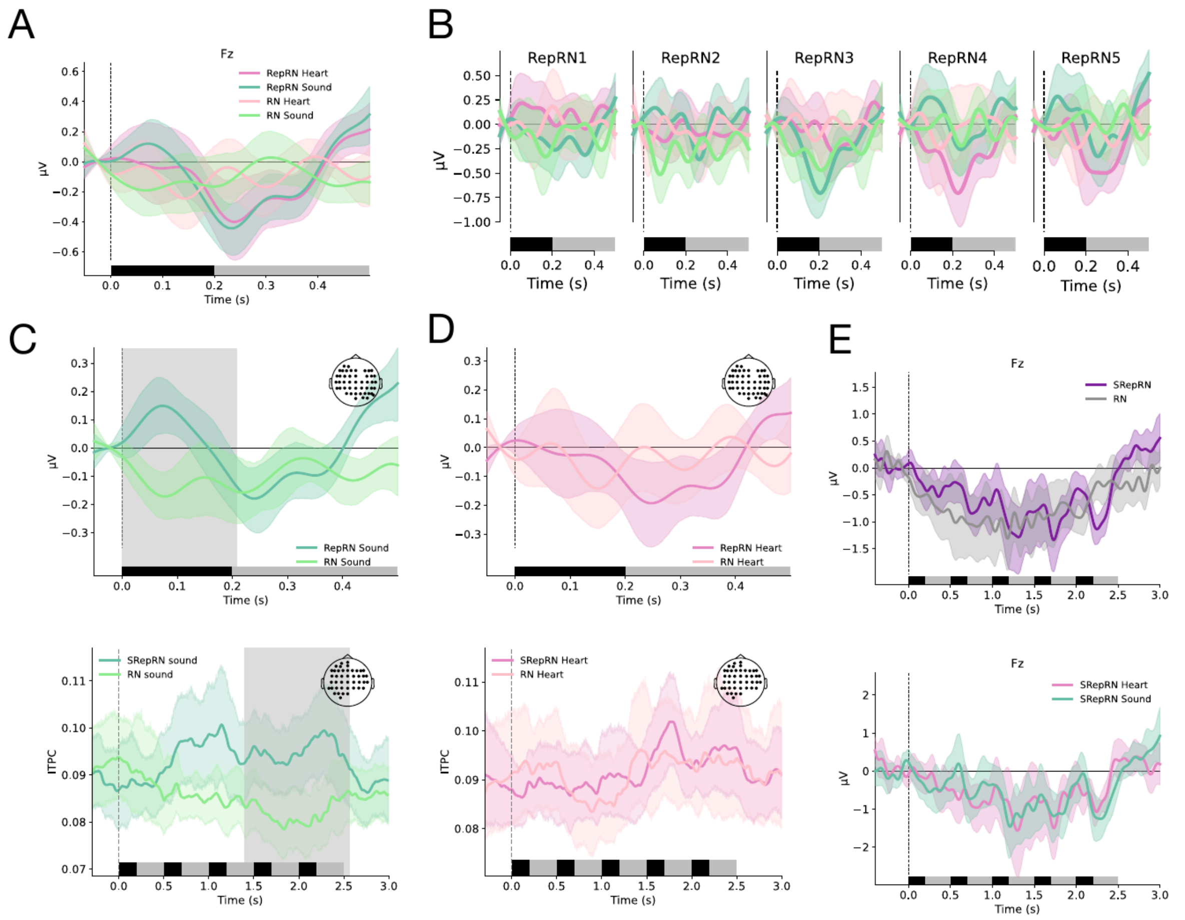

**SFig. 5**. Noise repetitions are better encoded during sound attention trials. **A**. Average brain response for all noise repetitions (RepRN) for Fz electrode. **B**. Average brain response for each of the five repetitions comprising the structured repetition (SRepRN) across channels shown in C. **C**. Top. Average response across channels that form the significant cluster for the difference between RepRN-Sound and RN-Sound. Bottom. Average inter-trial phase coherence in the delta band for RepRN-Sound and RN-Sound. Grey shading marks the temporal span of clusters and black dots indicate channels in clusters. The ERPs shown are the average voltage across channels for each cluster with a 95% bootstrap confidence interval. **D**. Top. Average response across the same channels as in C but for RepRN-Heart and RN-Heart. Bottom. Average inter-trial phase coherence in the delta band for the same channels as in C for RepRNHeart and RN-Heart conditions. **E**. Top. Average brain response for the structured repetition (SRepRN-Heart and SRepRN-Sound) in purple and for segments of random noise (RN-Heart, RN-Sound) in grey for Fz electrode. Bottom. Average

brain response for the structured repetition during attention to sound in green (SRepRN-Sound) and during attention to heart in pink (SRepRN-Heart) for Fz electrode. RepRN-Heart: random noise seeds presented during heart attention, RepRN-Sound: random noise seeds presented during sound attention, RN-Heart: plain white noise during heart attention trials, RN-Sound: plain white noise during sound attention trials.

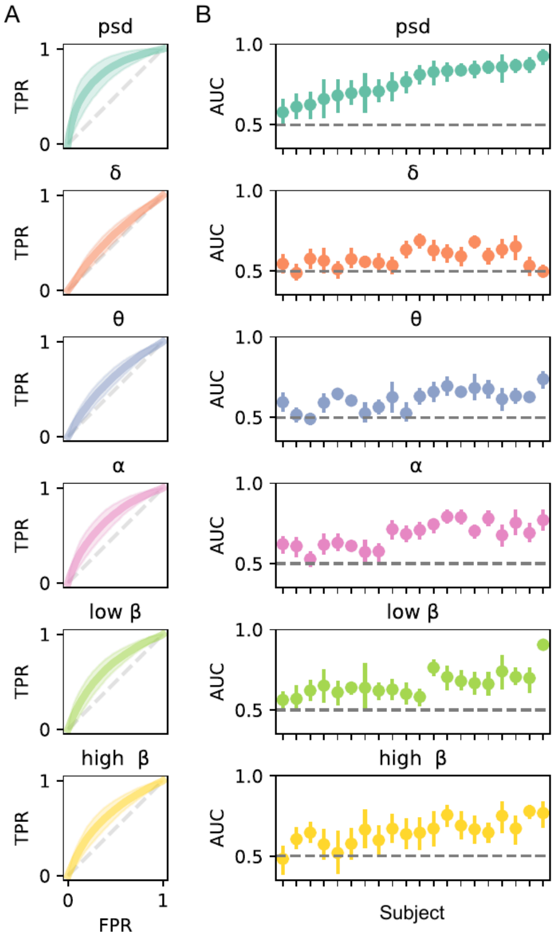


**SFig 6**. Power band-specific classifiers. Subject-level classifiers for each frequency band (delta, theta, alpha, low-beta, and high-beta). **A**. Average receiver operating characteristic curves across cross-validation folds and subjects. PSD: 0.75 ± 0.10, delta: 0.57 ± 0.06, theta: 0.60 ± 0.06, alpha: 0.67 ± 0.08, low-beta: 0.65 ± 0.07, high-beta: 0.64 ± 0.07. **B**. Average AUC across folds with 95% bootstrap confidence interval for each subject and classifier. Subjects are sorted considering the AUC for the classifier with all the frequency features (PSD).

*Adaptation for unresponsive patients*

Modifications were introduced to the original experiment to render the task more adequate for patients. The trial length was reduced to 18 seconds and delivered to patients in blocks of 28 trials separated by 2 min pauses. Each block was composed of 14 trials of heart-directed attention and 14 trials of sound-directed attention. Trial presentation was randomized within each block and a jitter of 1.5-2 s was introduced between trials. The AmN was 450 ms long and could appear between 1 and 3 times per trial, and the same 4 noise seeds were presented during both types of trials. Instructions were recorded in French and Italian and the task was presented via a custom-built Arduino stimulation box that sends the audio through Etymotic ER3C earphones and event markers directly to the amplifier. No report was asked after each trial and therefore no interoception or exteroception accuracies were computed. Paris patients were recorded with EGI 256 channels HydroCel GSN net and ECG activity was recorded using the PIB box. Signals were acquired with a Net Amps 300 EEG Amplifier from Electrical Geodesics, Inc, digitized at 256Hz. In Milan, patients were recorded with a 64-channel BrainAmp DC amplifier (Brain Products) including 60 EEG channels and 2 bipolar derivations for the ECG and EOG activity (sampling rate 1000Hz, low cutoff 0.1 Hz, high cutoff 250 Hz).

EEG processing differed in the following steps. ICA was performed on 1:30 Hz filtered data to remove movement and ocular artifacts. Component rejection was carried out on data filtered between 0.1:30 Hz before obtaining epochs for the HEP and the sub-epochs for classifier purposes. Each 18 s epoch was segmented into five 4 s epochs with an overlap of 3 s. ICA was performed on 1:7 Hz filtered data and the components were rejected from data filtered between 0.1:15 Hz, before epoching to obtain AmN (-0.1:0.9 s). Finally, data was combined into a 64-channel Biosemi layout configuration. For the classifiers involving the HEP, the canonical cluster obtained from the healthy participants' group analysis was used to select the features. Each classifier for each patient was run 100 times and the mean value across runs was compared to 1000 runs of surrogate classifiers. Finally, patient differences in the HEP were evaluated by averaging the EEG response to heartbeats across electrodes in the canonical clusters obtained for healthy participants. The averaged response was subjected to a point-by-point one-way ANOVA for time points -0.1:0.5 s, and Bonferroni corrected for multiple comparisons.

**Table S1. Demographics and behavioral assessment of patients**

| Code | Age at injury | Sex | Injury type | Days since injury | CRS-R | | | | | | | Diagnosis |
|---|---|---|---|---|---|---|---|---|---|---|---|---|
| | | | | | Au | V | M | OM | C | Ar | Total | |
| P1 | 61 | M | Hypoglycemia | 14 | 0 | 0 | 0 | 0 | 0 | 1 | 1 | VS/UWS |
| P2 | 40 | M | CA | 48 | 2 | 1 | 2 | 1 | 0 | 1 | 7 | VS/UWS |
| P3 | 22 | F | Encephalitis | 43 | 1 | 1 | 1 | 1 | 0 | 0 | 4 | VS/UWS |
| P4 | 38 | F | Brainstem stroke | 34 | 4 | 5 | 2 | 1 | 1 | 2 | 15 | LIS |
| M1 | 50 | M | TBI | 238 | 1 | 4 | 2 | 1 | 0 | 2 | 10 | MCS- |

Notes: P: patients assessed in Paris center, M: patients assessed in Milan center, CA: cardiac arrest, TBI: traumatic brain injury, Au: auditory functions, V: visual functions, M: motor functions, OM: oromotor/verbal functions, C: communication scale, Ar: arousal scale, VS/UWS: vegetative state/unresponsive wakefulness syndrome, LIS: locked-in syndrome, MCS-: minimally conscious state minus
